# MiFoDB, a workflow for microbial food metagenomic characterization, enables high-resolution analysis of fermented food microbial dynamics

**DOI:** 10.1101/2024.03.29.587370

**Authors:** Elisa B. Caffrey, Matthew R. Olm, Caroline Isabel Kothe, Joshua Evans, Justin L. Sonnenburg

**Affiliations:** Department of Microbiology and Immunology, Stanford University School of Medicine, Stanford, CA, USA; Chan Zuckerberg Biohub, San Francisco, CA, USA; Center for Human Microbiome Studies, Stanford University School of Medicine, Stanford, CA, USA; Sustainable Food Innovation Group, The Novo Nordisk Foundation Center for Biosustainability, Technical University of Denmark

## Abstract

**Summary:** Fermented foods, which contain a diverse array of microbial metabolites and microbes, are increasingly recognized as potential mediators of human immune and metabolic health. While there is growing interest in characterizing the microbial landscape of fermented foods, current characterization methods typically rely on 16S rRNA sequencing or marker gene-based methods, which have low taxonomic resolution and cannot functionally characterize microbes. Here we describe MiFoDB-workflow, a metagenomics workflow for the identification of microbes associated with food fermentation based on a primary database of 675 genomes of bacteria, yeast, fungi, and common fermented food substrates. We constructed the database using metagenome-assembled genomes (MAGs) derived from metagenomic sequencing of 90 fermented foods, combined with previously-published fermented food-derived MAGs, and relevant genomes deposited in RefSeq and GenBank genomes. We demonstrate the utility of MiFoDB for high confidence genome identification, including discovery of previously uncharacterized species, strain tracking across related foods, and functional analysis in fermented foods of different substrates. The workflow streamlines high-confidence characterization of the diversity of the fermented food landscape including novel ferments, and allows strain-level tracking of microbes across time, substrates, and producers.

---

Food fermentation has been used for thousands of years as a method of flavor enhancement, preservation and promoting food safety, involving the conversion of food components by microbes into desired food products^1^. Microbes associated with food fermentation are of growing interest for their ability to produce edible biomass and food ingredients for sustainability applications^2^. Fermented foods also modulate human health^3–5^, increasing gut microbiome diversity and decreasing markers of inflammation^6^.

Sequencing technology has allowed for the identification of fermented food-associated microbes that are difficult to isolate in culture. 16S and ITS sequencing provides a broad census of microbes present in a sample, while metagenomic sequencing provides genomic resolution of fermented food microbiomes^7,8^. Metagenomic sequencing enables resolving microbial functional attributes as well as species-and strain-level diversity of fermented foods, which can be particularly informative for spontaneous ferments, which typically harbor extensive microbial diversity and may be unpredictable^9^.

However, many challenges remain in taxonomic profiling of fermented foods. The most commonly used methods today rely on clade-specific marker genes (e.g., MetaPhlAn^10^) or k-mer-based profiling (e.g., Kraken^11^, Clark^11,12^ Kaiju^13^). These methods are computationally efficient, but the programs implementing these approaches generally have the following shortcomings: 1) report relative abundance values scaled to 100%, preventing assessment of the amount of unclassified reads present in a sample, 2) cannot report metrics like breadth of coverage to assess the confidence that a particular microbe is detected 3) do not provide assessment of microbial genes detected, and 4) in practice are almost exclusively used with their provided reference databases, precluding detection of novel genomes. The MiFoDB-based workflow developed and described here overcomes all these shortcomings by leveraging full genome characterization, providing high accuracy in genome detection, ability to detect novel genomes, strain level resolution and tracking, and functional insight into microbial communities.

Profiling the composition and function of microbial consortia using metagenomic sequencing is aided by a high-quality reference database of relevant genomes. Such databases exist for the human gut^14^, soil^15^, ocean^16^ and lake^17^. However, no such resource is available for fermented foods. Other databases for fermented food sequence analysis are not compatible with large-scale metagenomic data and are not suitable for identification of novel MAGs^18^. Here we present MiFoDB (Microbial Food DataBase), a workflow for the identification of bacterial and eukaryotic genomes in fermented foods, with an accompanying customizable database. The database integrates novel genomes (MAGs) derived from newly reported metagenomic data generated from a variety of fermented foods plus previously published MAGs^7,18–24^, along with RefSeq and GenBank genomes. Each unique species is represented by the highest quality reference genome. MiFoDB includes 586 medium-to high-quality unique prokaryotic reference genomes across 9 phyla, including 45 previously unnamed species and 82 unique eukaryotes reference genomes, including 4 potential novel species. In addition, incorporation of 7 common fermented food substrate genomes allows for more precise mapping of metagenomic sequences. Application of the full MiFoDB-based metagenomic workflow easily allows other users to conduct strain-resolved, functional analysis of their fermented food metagenomes.

We characterized 90 distinct samples from 56 fermented foods, which include four ferments sampled at multiple timepoints, and identified a total of 779 high-confidence genomes across samples. Downstream analyses include strain-tracking^25^, where identical microbial (>99.999% ANI) strains were traced across fermented foods, passages (sequential ferments), timepoints, and batches. Correlating strain abundance with metabolomics results from a semi-targeted microbiota metabolite reference library^26^ identified strain specific metabolite signatures, including production of histamine, a molecule with health relevance. Finally, functional analysis showed differential expression of CRISPR-associated domains, CAZymes, and Multi-Drug Resistance (MDR) genes in vegetable and dairy ferments.

As interest in food fermentation grows, gaining a high-resolution and comprehensive view of microbes involved in distinct fermentation processes will impact biomedicine, food safety, and food quality^1,27^. This full MiFoDB-based metagenomic workflow easily allows other users to conduct strain-resolved, functional analysis of their fermented food metagenomes. A full description of the MiFoDB workflow and accompanying open-source database is available in GitHub (https://github.com/elisacaffrey/MiFoDB) and documentation is available online (https://mifodb.readthedocs.io/en/latest/).

## Results

### Metagenomic sequencing of a diverse collection of fermented food samples

Characterizing the microbial landscape of ferments remains an important challenge to understand which microbes play a role in fermentation and how they interact within their community to influence food safety, flavor and human health. Time-course metagenomics gives insight into community changes during fermentation, identifying microbes that drive community composition which might not be identified at completion of fermentation. In addition, while metagenomic research has primarily focused on dairy ferments, non-dairy ferments including soy and vegetable fermented foods remain underrepresented.

We performed metagenomic sequencing on 56 fermented foods collected between January 2021 and March 2023 (Supplementary Table 1). From these 56 ferments, the majority of which were vegetable ferments, we sequenced 90 samples in total including across various time points and from different parts of the food (Fig. 1a). Fermented foods with time-series samples collection include two green cabbage sauerkraut (6 samples), red sauerkraut (3 samples), goat kefir fermentation (16 samples), viili (2 samples) and tempeh (2 samples). To gain a deeper understanding of microbial localization in a fermented food, the brine and cabbage samples of kimchi (4 total, including 2 different timepoints), red sauerkraut (2 samples), and the supernatant and sediment of wine and cider (4 ferments, 9 samples) were sequenced. Samples from only the final timepoints of soy-based ferments, vegetable ferments including sauerkraut and kimchi, wines, and cheese were included as well. DNA extraction was performed using ZymoBIOTICS DNA Miniprep kit, and subjected to paired-end Illumina shotgun sequencing (Fig 1c). A total of 776.56 Gbp of data was generated from all 90 samples (range = 0.12–26.14 Gbp; avg = 8.628497 ± 0.63 Gbp). Following pre-processing, assembly, and binning, we recovered a total of 1,186 microbial genomes (see methods for details).

**Figure 1:**
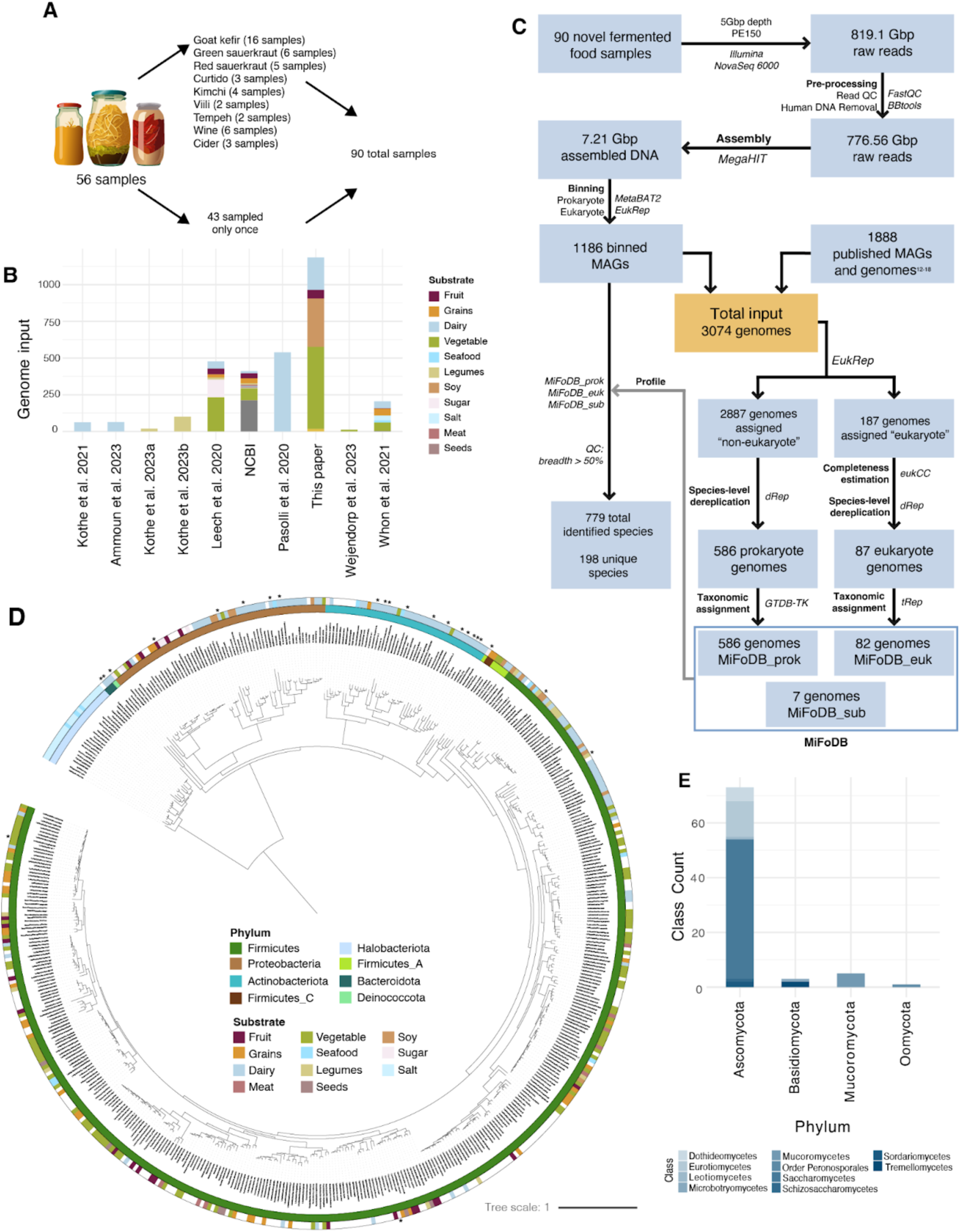
MiFoDB workflow and genome database generation. A. Overview of sample collection for shotgun metagenomic sequencing of fermented food samples. B. Summary of all input genomes by each genome substrate origin, including MAGs assembled in this paper, previously published MAGs and reference genome. C. Workflow for generation of genomes included in MiFoDB.. D. A phylogenetic tree of prokaryotic genomes included in the final MiFoDB_prok reference. The tree is decorated by phylum (inner ring) and substrate of origin for each reference genome (outer ring). E. Frequency of the eukaryotic genomes included in the final MiFoDB_euk reference by phylum; shade indicates class. A member of order Peronosporales does not belong to an assigned class, so is listed by order.

### Defining a broad and representative collection of microbes associated with fermented foods

Recognizing how common, non-fermented food targeted reference databases may miss microbes, we proceeded with the creation of a novel, custom fermented-food focused metagenomic database. To perform high-confidence profiling of the recovered MAGs and for ease of downstream diversity, taxonomic, and functional analyses, we first collected genomes associated with fermented foods. MAGs assembled from the 90 samples were collated with genomes from previous fermented food sequencing to create a high quality reference database of relevant genomes. These sources include the ODFM database^18^ (205 genomes), three dairy ferment metagenomic surveys^20,21,28^ (539, 62, and 64 respectively), a large metagenomic survey of substrate-associated differences in fermented food MAGs^19^ (477 genomes), and three studies of novel fermented foods^24,23,22^ (19, 100, 11 genomes respectively). Due to the central role members of the *Lactobacillus* genus play in lactic acid fermentation, 320 *Lactobacillus* NCBI reference genomes reported in the literature were included^29^. In addition, the reference genome from microbes found in kombucha and wine-associated yeasts were added from NCBI^30^ (18 and 73 genomes respectively).

The 3074 genomes in the input set reflect a wide global distribution of sample origin from which genomes were derived (Supplementary Fig. 1a), with overrepresentation of samples from the United States due to the large contribution of novel MAGs from our own sequencing to the database (Supplementary Table 1). MAGs and reference genomes originating from dairy and vegetable ferment dominated the total input set (Fig. 1b; Fig. 1c, orange box). Overall, this genome set represents an up-to-date, global sampling of fermented food microbes.

### Generation of the MiFoDB genome database

A primary goal of establishing this database of fermented food microbial genomes is to enable rigorous metagenomic analyses, including taxonomic profiling, strain-tracking, and functional annotation. The presence of multiple closely-related genomes in the same database is known to thwart these efforts^25^. Thus, we next selected representative genomes for each species-level group present in our input set (Fig. 1c, orange box). We opted for a tripartite database strategy, featuring MiFoDB_prok for prokaryote (bacteria and archaea) identification, MiFoDB_euk for eukaryote identification, and MiFoDB_sub for substrate identification (Fig. 1c). This modular approach allows for the use of domain-specific thresholds, higher accuracy in gene calling and targeted downstream analyses. We implemented a custom genome dereplication pipeline for determining which genome of a species would be the representative (see methods for details), and ultimately identified 586, 82, and 7 species-level representative genomes in the prokaryotic, eukaryotic, and substrate databases, respectively (Supplementary Table 2).

The final MiFoDB_prok database contains 586 genomes, representing 558 bacterial and 28 archaeal genomes (Fig. 1d; Supplementary Fig. 1b), with 505 genomes (over 86%) at >90% completeness and <5% contamination (Supplementary Fig. 1c). Taxonomy was assigned to any MAGs in MiFoDB_prok using GTDB (Genome Taxonomy Database)^31^, while tRep (https://github.com/MrOlm/tRep/tree/master/bin) was used to assign taxonomy to any MAGs in MiFoDB_euk. Any genomes originally added from NCBI maintained their assigned taxonomy. The majority of genomes serving as representatives for each species came from NCBI (318/586 genomes), reflecting the higher genome quality of deposited isolated reference genomes compared to de novo assembled MAGs (Supplementary Fig. 1d). Bacterial genomes within the DB are dominated by Firmicutes, primarily of genus *Lactobacillus*, reflecting the dominance of the lactic acid microbe reference genomes. While all archaeal genomes were identified in sequenced salt used in kimchi making or seafood ferments, bacteria reflected a wider distribution of substrates (Fig. 1d). Of note, no taxonomic identification was assigned at a species-level to 45 bacterial genomes, which were only identified at the genus level (Supplementary Fig. 2a, Supplementary Table 3). These novel species are members of *Acetobacter, Arthrobacter_H, BOG-930, Canibacter, Chromohalobacter, Companilactobacillus, Enterobacter_D, Ethanoligenens, Galactobacter, Gluconobacter, Halomonas, Idiomarina, Korarchaeia, Leuconostoc, Liquorilactobacillus, Luteimonas, Microbacterium, Nocardiopsis, Novosphingobium, Pantoea, Proteus, Pseudoalteromonas, Psychrobacter, Psychroflexus, Rothia, Salinisphaera, Savagea, Sphingobacterium, Streptococcus, Streptomyces, Tatumella, Tetragenococcus, Yaniella,* and two species in each of *Brachybacterium, Brevibacterium, Corynebacterium, Halomonas, Lactiplantibacillus, Levilactobacillus.* Two prokaryotic species were only identified at higher taxonomic levels, including a member of family Micrococcaceae and an archaeal member of class Korarchaeia. Novel MAGs originated from 8 different countries, including Belgium, Brazil, Denmark, Germany, Ghana, Ireland, the United States and the United Kingdom (Supplementary Fig. 2b-d). While identification of novel MAGs from underrepresented countries was not surprising, identification of novel MAGs from larger dataset from the United States and Ireland highlight just how understudied fermented food microbiomes remain. Expansion of this database is expected with future sequencing of novel samples from diverse geographic regions.

The final MiFoDB_euk database contains 82 representative eukaryotic genomes. Following taxonomic identification of these genomes, 5 likely represented substrate genomes, rather than microbial eukaryotes involved in fermentation. Specifically, we assembled two genomes from genus *Bos* (cattle), and one genome of each: *Brassica oleracea* (wild cabbage), *Oryza sativa* (rice), and *Brassica rapa* (field mustard) genome. Interestingly, completeness scores for all these bins was >50%, with the highest at 84.4% completeness (Supplementary Table 4). As these substrate genomes do not represent microbes involved in fermentation, these bins were removed from the MiFoDB_euk database, resulting in the 82 eukaryote members in the current MiFoDB_euk (Supplementary Table 5, Fig. 1e). These eukaryotic microbial genomes include four annotated as *Kazachstania saulgeensis* but with >95% ANI, indicating these are likely to represent 3 or more novel close relatives to *Kazachstania saulgeensis*. *Plasmopara viticola*, a fungus-like eukaryote of phyla Oomycete, is the only eukaryote in the database without an assigned class.

Finally, for identification of substrate genomes in the metagenome sample, MiFoDB_sub was developed. Briefly, the database consists of 7 high-quality RefSeq genomes from *Brassica oleracea var. oleracea* (wild cabbage plants*), Bos taurus* (cow), *Capra hircus* (goat), *Vitis vinifera* (wine grapes), *Glycine max* (soybean), *Oryza sativa* (rice), and *Triticum aestivum* (common wheat) (Supplementary Table 6).

### MiFoDB allows for detailed mapping insight

We wanted to apply the tripartite MiFoDB database (prokaryotic, eukaryotic, substrate) to profile all fermented food metagenomes sequenced in this study (see methods for profiling details). The MiFoDB database recruited the majority of reads in most samples (mean = 61.4% ± 2.06% of reads) (Fig. 2a). However, considering the typically lower absolute mapping success of samples^35,36^, fermented food sample mapping reflects an expected middle ground in diversity, being less diverse than the soil, but underexplored compared to the gut microbiome. In a few samples, the substrate genome mapped up to 91% of the total sequenced genome. Inclusion of substrate reference genomes to the MiFoDB workflow allows for a deeper understanding of the fermented food profile, suggesting that use of bioinformatics methods that artificially sum all relative abundance values to equal 100% (including MetaPhlAn and Kraken) greatly skews our understanding of microbial diversity in the sample (Supplementary Fig. 3a).

**Figure 2.**
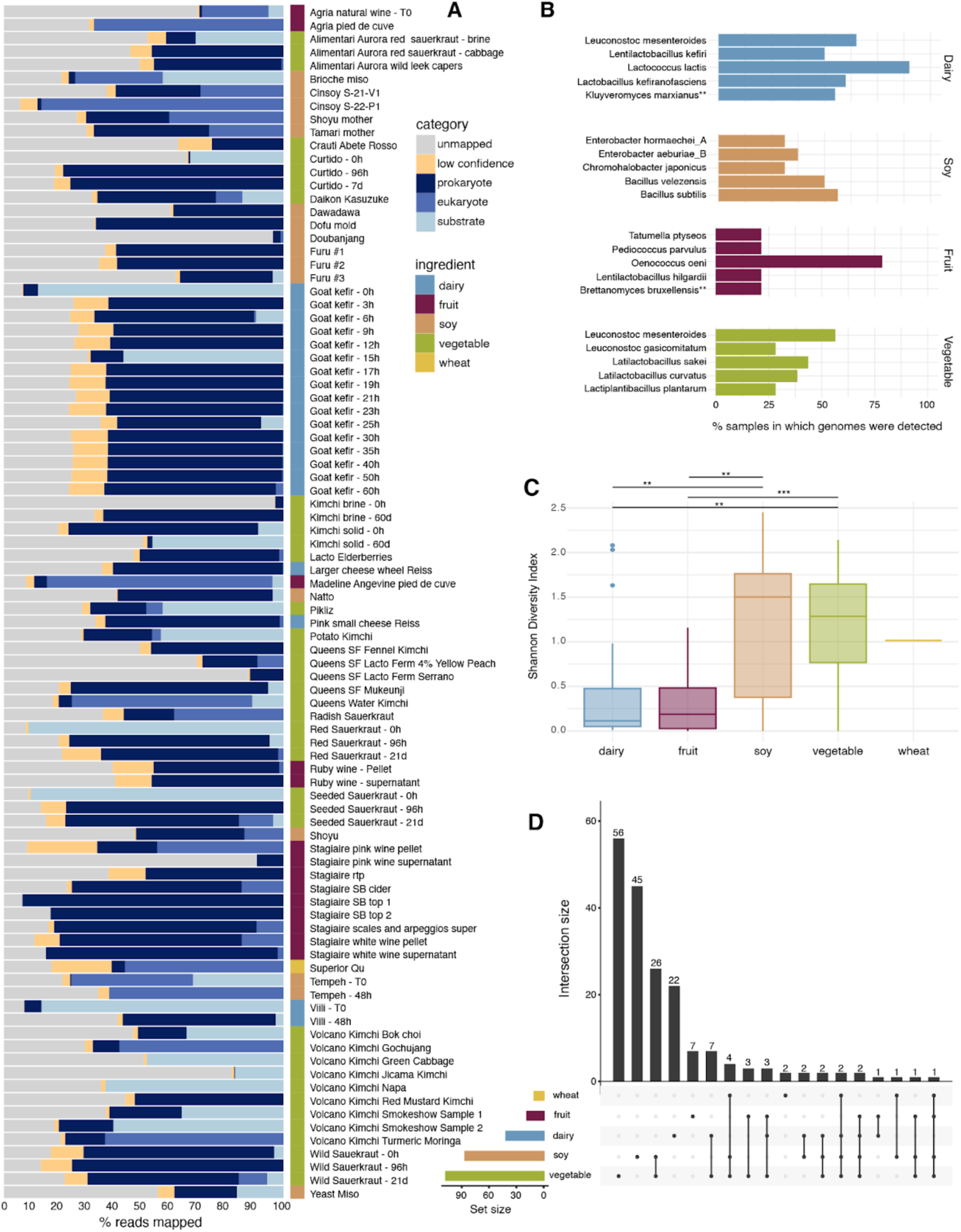
A. Profiling results of fermented food samples using MiFoDB, mapped to prokaryotic, eukaryotic, or substrate. Low confidence profiling results (breadth <0.5) and unmapped reads are also noted. B. Bar plot of the five most frequently identified microbes by substrate category. Eukaryotes are differentiated with (**). Wheat samples are not included, as only one wheat-based ferment was sequenced. C. Shannon diversity index of microbes (eukaryotic and prokaryotic) profiled by ingredient category. p-values from Tukey’s test; p < 0.05 (**), p<0.001 (***). D. UpSet plot of intersecting species across ingredient types. The bar plot above represents the number of total species within each set. The rows below each represent an ingredient, with the edges connecting nodes that are included in the set. The bar plot on the left shows the total number of species for each substrate.

An important consideration in profiling metagenomic data is the presence of unmapped reads. Species with completely novel genomes, such as the 45 mentioned above (i.e., with previously unidentified genomes) present in a given ferment are one source of unmapped reads; such novelty requires MAG assembly and addition to the database prior to mapping reads. A second source of unmapped reads could be due to species with known genomes (i.e., in GTDB) that are currently not in our database but are relevant to different fermented foods. To address this second scenario as a basis of unmapped reads we profiled all our samples against GTDB, the Genome Taxonomy Database^31^, using sylph^37^, a novel metagenomic profiling program. Profiling resulted in the reported identification of 1068 unique microbial genomes, 603 of which were not included in MiFoDB. Reference genomes from microbes identified in more than one sample or with a reported abundance > 0.5% were incorporated in a new version of MiFoDB (for a total of 338 additional genomes) and our 90 samples were profiled against it (Supplementary Fig. 3b). The majority (76%) of added genomes reported a total abundance across all samples of < 1%, with the total mean of mapped reads increasing only slightly from 61.4% ± 2.06% to 62.9% ± 2.08%. Due to the small change in abundance of mapped reads, and that the majority of newly added genomes mapping >1% were only identified in one sample each, we chose to not add these to the database. However, use of sylph to identify and incorporate novel genomes to the core database could aid in the mapping of understudied and novel ferments.

MiFoDB reflects the expected microbial content of fermented foods for the different substrates (Fig. 2b)^19,38–43^. In the dairy samples, L. lactis was present in 90% of dairy-based samples, along with *L. mesenteroides, L. kefiranofaciens, L. kefiri*, and the yeast *K. marxianus* reflecting the representation of kefir samples in the data. Soy and vegetable substrate ferments had higher diversity (Fig. 2c), with *B. subtilis* and *L. mesenteroides* present in just over 50% of samples, respectively. *O. oeni*, a key player in malolactic fermentation, was observed in >78% of fruit (wine) samples, reflecting the dominance of the microbe at the final stages of wine production. Comparing presence of microbes across substrates, vegetable ferments had the greatest number of unique genomes (Fig. 2d). No species was identified in all five ingredient categories; however, this might be a reflection of the limited number of samples for certain ingredients (wheat, fruit) rather than a lack of overlapping microbes.

Most microbes contributing reference genomes were originally isolated or identified in food sources, as reported in literature searches and existing sample database information (see Supplementary Table 2 for sources of microbes contributing genomes to the database). However, several species identified in our fermented foods have a reference genome isolated from non-food sources. These include: *Levilactobacillus brevis* isolated from feces, *Kodamaea ohmeri* isolated from honeybee, *Weissella cibaria* isolated from infant saliva, *Rhizopus oryzae* isolated from tobacco, and *Secundilactobacillus silagei*, *Pediococcus parvulus,* and *Loigolactobacillus coryniformis* isolated from silage. Based on relatedness to known fermentation-relevant microbes, and these existing reports of association with non-human food fermentation-practices (e.g., silage), supports the validity of these identifications. This identified novelty also supports our decision to include genomes in the database from groups of fermentation-adjacent microbes that have not yet been formally associated with fermented food.

To evaluate MiFoDB’s ability to characterize novel fermented samples, we took a closer look at superior qu, a poorly characterized wheat-based ferment used as a starter for alcoholic beverages produced through the wild fermentation of wheat berries. To highlight the importance of a microbial food specific database, profiling with inStrain was performed using either UHGG^14^, the Unified Human Gastrointestinal Genome, or MiFoDB. Profiling reads from superior qu using either of these two databases revealed the better performance of MiFoDB over UHGG (61.2% high-confidence mapped reads vs. 3.12%; Fig. 3a).

**Figure 3:**
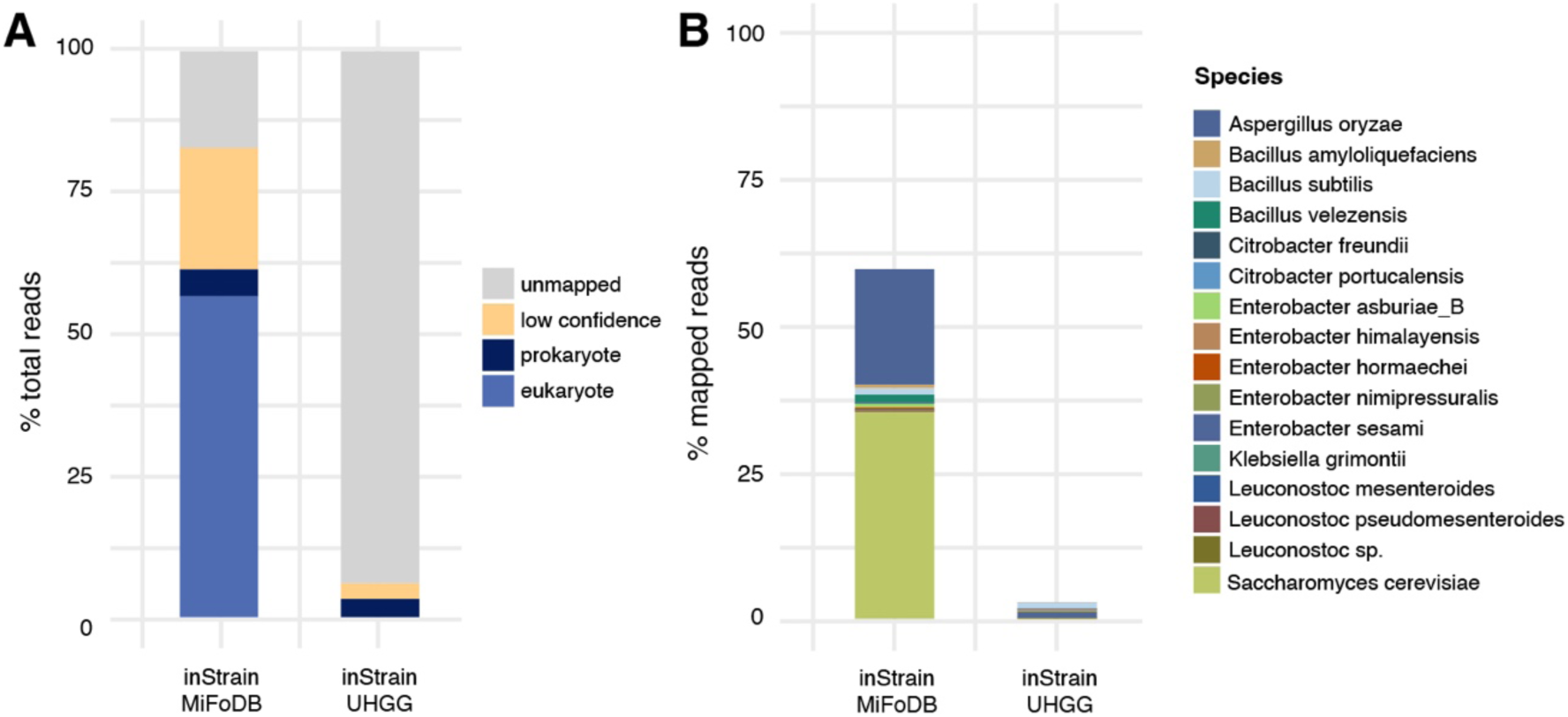
Comparison of alignment-based profiling databases. A. Comparison of percentage of reads mapped against MiFoDB versus UHGG for the novel sample, superior qu. Mapped reads include identified prokaryotes and eukaryotes, as well as low confidence reads mapped (breadth <0.5) and unmapped reads. 61.2% of reads mapped with high Confidence to MiFoDB versus 3.12% for UHGG. B. Abudance of high confidence mapped reads (breadth>0.5) using both MiFoDB and UHGG reference databases. Using a fermented food focused database increases read mapping by >58%.

Comparing species identified using UHGG or MiFoDB (Fig. 3b) revealed that UHGG lacked two of the most abundant eukaryotes identified using MiFoDB, *Aspergillus oryzae* and *Saccharomyces cerevisiae*. Of note, 20.3% of the absolute abundance of the superior qu sample was reported as the filamentous fungus *Aspergillus oryzae*. *A. oryzae* is closely related to *A. flavus*, a pathogenic fungus known for aflatoxin production^44^. While accurately differentiating between *A. oryzae* and its aflatoxin producing relatives is of clear importance for food safety, *A. oryzae* and *A. flavus* share an ANI of 0.992245 (Supplementary Fig. 4, Supplementary Table 7) making computational differentiation a challenge. Due to the high ANI, *A. oryzae* and *A. flavus* cannot be differentiated using MiFoDB, and only *A. oryzae* is included. However, additional analyses indicated presence of *A. oryzae* and not *A. flavus* in our samples (Supplementary Note 1). Out of abundance of caution any samples that map to *A. oryzae* should not be assumed as aflatoxin-free without further confirmation.

### Strain tracking across samples

An important advantage of assembly-based genome profiling is the ability for microdiversity-aware genome comparison and strain identification. By profiling our samples with inStrain^25^ MiFoDB can be used for unbiased identification of high-confidence strain shared across samples (Fig. 4a, Supplementary Table 8), including strains shared through passaging, in ferments made by the same producer, and across timepoints. In tamari and soy sauce fermentation strains of *Bacillus velezensis* 202_3 and *Ligilactobacillus acidipiscis* 187_1 were identified in both the shoyu and tamari starters (termed “mothers”), as well as the ferment produced after adding the starter (Fig. 4b). Strains can also be identified across ferments made at different time points by the same producer (Fig. 4c). Three samples of the same cider, a pinot noir rosé, and a sauvignon blanc and chardonnay blend, with fruit deriving from different geographic areas and produced across the span of a few months shared the same strain of *Oenococcus oeni* 184_1, likely derived from the shared fermentation environment. To benchmark our results, we compared inStrain to StrainPhlan4^45^, a strain tracking method based on consensus ANI measurements that requires a relational-based input. We inferred false-positives when strain sharing was called between samples of different provenance (making strain sharing exceedingly unlikely). While inStrain resulted in a 2.2% false positive rate (3/137 strain sharing events; two events were between samples collected and sequenced at the same time, suggesting technical/physical contamination rather than bioinformatic error), StrainPhlan4 resulted in 12.5% false-positive rate (17/136 strain sharing events between unrelated samples) (Supplementary Tables 8-9).

**Figure 4:**
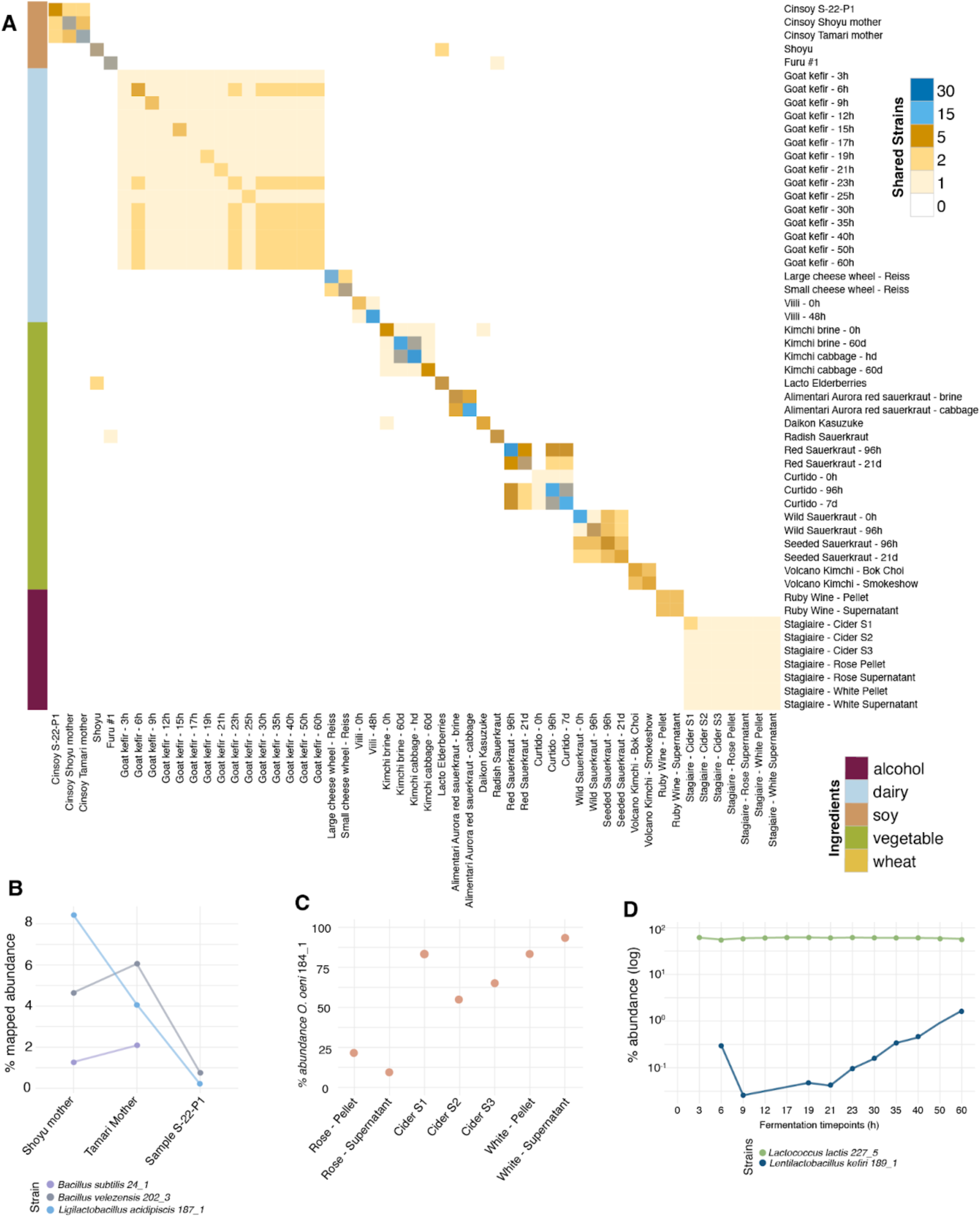
Strain sharing across fermented food samples. A. inStrain-determined strain sharing between samples. Strains are primarily shared between related samples (same samples across different timepoints, samples made by the same producer, and samples used as a starter for a second sample). B. Strains tracked between starters (Shoyu mother, Tamari mother) and packaged-product (Sample S-22-P1). C. One strain of *Oenococcus oeni* tracked across samples of cider and wine made without added starters by the same producer at different time points. D. Two strains tracked in goat milk kefir across 14 timepoints.

During milk kefir fermentation, strain *Lactococcus lactis* 227_5, likely present but below the limit of sequencing detection at 0 hour time point, reached close to 50% absolute abundance in the sample, and was easily identified at all following timepoints, remaining at a similar abundance across the 60 hour fermentation period. *Lentilactobacillus kefiri* 189_1, not identified before 6 hours, remained at lower abundance, but was still traced across the ferment, in increasing abundance after 23 hours to the final fermentation time point (Fig. 4d).

Production of metabolites within microbial communities plays a role in structuring metabolic interactions, and when occurring in fermented food, can impact food safety, shelf life, flavor, and potential health properties. Characterizing metabolite profile at a strain level has the potential to aid selection and use of specific strains in food production. Coupling semi-targeted metabolomics of a milk kefir time-course series with metagenomics strain data, we observe strain specific metabolite signatures (Supplementary Fig. 5). *L. kefiri* 189_1, a heterofermenter, increases steadily over the course of fermentation. Abundance correlates to higher abundance of organic acids and their derivatives, like proline and creatine. Metabolites correlating to *L. lactis* 227_5, a homofermenter which increased rapidly in the initial three hours of fermentation following inoculation of the milk substrate with the kefir grains do not show as strong of a correlation as *L. kefiri* 189_1, likely due to the difference in microbial abundance over the course of fermentation time. Multi-omics application of MiFoDB to strain-level metabolite profiles allows for a deeper understanding of the interrogation of strain-specific microbe-substrate interactions, and has important implications for food safety, taste profiling, and customization of fermented products. For example, correlation of *L. kefiri* 189_1 abundance to histamine production highlights the need to better characterize *L. kefiri* histamine production and identify potential regulatory mechanisms, at a strain, species, or microbial community level, to mitigate histamine production and to create a kefir that is appropriate for those with bioamine sensitivities.

### MiFoDB allows for robust functional profiling

A key advantage of our MiFoDB workflow is the ability to directly profile microbial functional potential. To explore the use of MiFoDB in functional profiling of fermented foods, genes identified from our fermented samples were annotated against Pfam^46^ (protein family), CAZy^47^ database (carbohydrate-active enzymes (CAZymes)), and CARD^48^ (antibiotic resistance genes) (Supplementary Table 10). Comparing fermented dairy and vegetable ferments, CRISPR-associated domains (Cas1, Csn2, Cas9 PI, Cas2, HNH4) were enriched in vegetable ferments, while dairy-ferments were significantly associated with phage-related genes (L_lac_phage_MSP, Lac_bphage_repr, Phage_Treg, TerB-C, TerB-N, CW_7) and bacteriocin expression (Lactococcin, Helveticin-J). Interestingly, Pectate lyase 4, an enzyme family in plants involved in maceration and rotting of plant tissue, is significantly associated with microbial vegetable fermentation while depleted in dairy fermentation (Fig. 5a).

**Figure 5:**
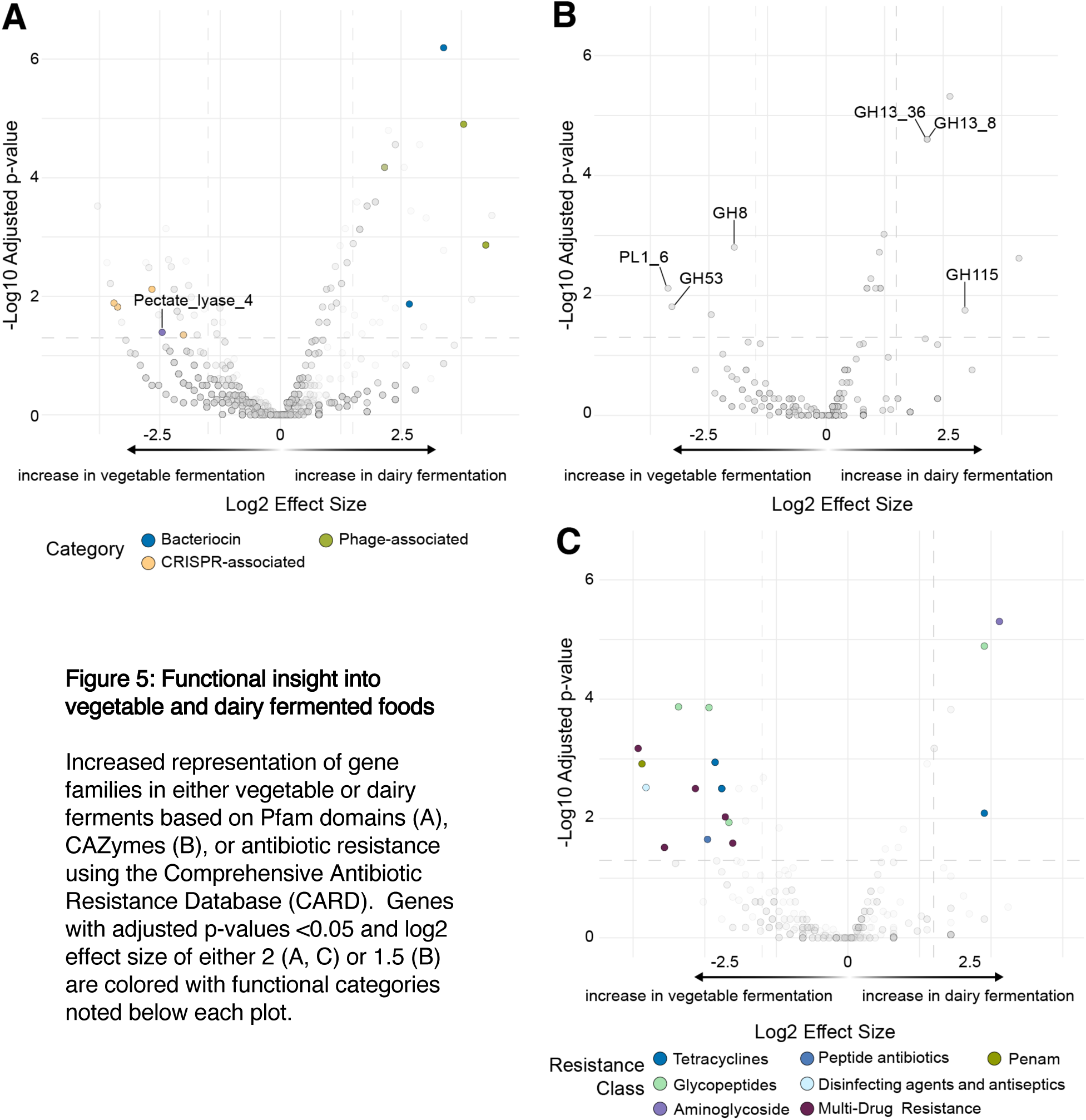
Functional insight into vegetable and dairy fermented foods. Increased representation of gene families in either vegetable or dairy ferments based on Pfam domains (A), CAZymes (B), or antibiotic resistance using the Comprehensive Antibiotic Resistance Database (CARD). Genes with adjusted p-values <0.05 and log2 effect size of either 2 (A, C) or 1.5 (B) are colored with functional categories noted below each plot.

CAZy identified Glycoside Hydrolase Family 8 (GH8), formerly known as a member of the cellulase family, as enriched in vegetable ferments. In addition, vegetable ferments show enrichment in pectate lyase (PL1_6), and the enzyme GH53, an enzyme with specific activity on type I arabinogalactans, carbohydrate components of the primary cell walls of dicots^49^. Dairy fermentation was enriched for amylases (GH13_8 and GH13_36), and Glycoside Hydrolase Family 115 (GH115) which includes α-glucuronidase activity (Fig. 5b). Profiling against the CARD database (Fig. 5c) identified a number of antibiotic-resistant annotations, with vegetable ferments enriched for Multi-Drug Resistance (MDR) genes. Antibiotic resistance, as a naturally occuring defense in the warfare typical between members of competitive microbial communities, is not unexpected and has been previously discussed in fermented foods, particularly in lactic acid bacteria^19,50–52^. However, drug-resistance acquisition and impact on health from fermented food-associated microbes requires further study and contextualization with other microbial communities such as those from the human gut microbiome.

## Discussion

As scientific inquiry of fermented food continues to grow, appropriate tools are needed to characterize complex interactions of microbial communities. This study offers a workflow for the identification of MAGs from fermented foods, coupled with MiFoDB, a novel resource for the taxonomic profiling and functional analyses of ferments. This powerful tool to interpret sequencing data associated with fermented food samples can be applied to single time points or to monitor community dynamics across a time course or storage. Its tripartite system allows for the maximum flexibility in trans-kingdom identification and genome profiling. With over 600 medium to high-quality genomes of bacteria, archaea, eukaryotes, and common fermented substrates assembled from fermented foods or reported as having high relevance to fermentation, our database and approach allow for high-confidence identification of microbes shaping the ferment landscape.

Integrating MiFoDB into the alignment based taxonomic profiling program inStrain offers additional advantages. Reporting of mapped abundance rather than the relative abundance gives a better understanding of how thoroughly the fermentation landscape was characterized. Results easily integrate with functional analysis tools to elucidate KOfams or CAZymes, which inform our understanding of functional attributes including metabolic activities and mediators of interactions within the fermented food landscape. Coupling of functional analysis with metabolomics, particularly at a strain-specific level, allows for an even deeper understanding of strain-specific and community-level metabolic action. Metagenomic functional annotations and metabolomics aid in known compound identification and novel compound discovery. Coupling strains to the production of bioactive nutrients, antimicrobial compounds, and health-modulating metabolites is a key to informing high quality and reproducible fermentation practices including rational design and control of desired features within fermented foods.

Most interesting is the application of MiFoDB for precise strain tracking. In conjunction with inStrain, strains at a shared ANI of 99.999% (representing less than 10 years of evolutionary distance) can be tracked across fermented food samples. Strains can be traced with high-confidence across environments and timepoints, addressing questions of strain origin, evolution, and microbe-host relationship, including microbial transfer and engraftment into and interaction with our host microbiomes. This MiFoDB-workflow can help address interesting microbial ecology questions, like where in the environment microbes involved in spontaneous fermentation come from. Findings have important implications for cultural questions including, how do strains from a specific fermented food producer influence the final taste of the product, or how resilient are community members to strain invasion or phage infection?

Finally, identification of novel microbes and deeper characterization of understudied microbes involved in fermentation would be an important contributor to sustainability efforts in a number of fields^2^. In gastronomy, microbial novelty has the potential of encouraging flavors and textures on sustainable substates, either by applying microbes to novel substrates, or by applying principles of microbial ecology to promote specific culinary phenotypes of interest. Using microbes in production of animal feed has important potential in enhancing nutritional value of current underused nutritional sources. Novelty in metabolism and metabolite production of these microbes can add to the sustainability toolkit, potentially improving biofuel efficiency and materials needed for sustainable energy production.

The database presented here serves as a starting point for more comprehensive mapping of the tremendous global diversity of fermented foods. The current database primarily includes MAGs from foods originating in the United States, reflecting the current sample and resource availability for metagenomic sequencing. However, the flexible design allows for continued incorporation of novel MAGs and reference genome as the fermented food metagenomics field expands. As more metagenomes are added to MiFoDB, we will gain insight into the fermented food microbial landscape, with the goal of building a global genome collection.

This workflow is an important tool in ferment customization. Modifications to a specific strain or community could promote or prevent production of specific characteristics, like favorable-aromas and lactic acid, or off-flavors and bioamines. More broadly, application of MiFoDB to both novel and previously well-characterized fermented foods will help illustrate how technique and ingredients influence the microbial community. Detailed microbial and molecular characterization of fermented foods creates an avenue to understand the basis of how fermentation practices, such as modern and traditional methods, influence health and flavor.

In summary, MiFoDB offers a high-quality, comprehensive, and flexible metagenome reference database for fermented food metagenomic samples. As the field continues to expand, identification of novel microbes will enrich our understanding of microbial ecology and community development. The flexibility of this database and workflow outlines novel genomes and enables incorporation into the database, allowing for customization and expansion as the field continues to grow. In addition, the facilitated incorporation of strain tracking and gene profiling creates a powerful platform for the exploration of future work across numerous fields, including host-microbe interactions, food safety, culinary practice, and human health.

## Supporting information

High resolution figures

Supplementary Table 1: List of all samples

Supplementary Table 2: List of all genomes in MiFoDB_v2

Supplementary Table 3: Putative novel prokaryotes

Supplementary Table 4: Assembled substrate genomes

Supplementary Table 5: Eukaryote genomes in MiFoDB_euk

Supplementary Table 6: Substrate genomes

Supplementary Table 7: Aspergillus alignment results

Supplementary Table 8: inStrain results

Supplementary Table 9: StrainPhlan4 results

Supplementary Table 10: Functional analysis results

Supplementary Table 11: Kefir untargeted metabolomics

## End notes

### Acknowledgements

We are most thankful to all fermented food sample producers and contributors, including Dario Barbone of Alimentari Aurora, Sandor Katz, Mara King, Macklin Casnoff, Hannah Wastyk, Laura Krum, Aruna Lee of Volcano Kimchi, Clara Lee of Queens SF, Brent Mayeaux of Stagiaire Wine, Cody Reiss, Natalie Schrauwen, and Richard Shover. Thank you to Claire Clayton, Danny Ramos, Will Tospern and Adam Strozyk of No Lords Wine, and Alex Ma, Sarina Uriza Dito, Connor Geraghty, and Stephen Amato-Salvatierra of Ruby Wine. Thank you to Sailendharan Sudakaran for his advice on the analysis.

This work was supported by grants from the Bill and Melinda Gates Foundation, the NIH (R01-DK085025 (JLS), F32-DK128865 (MRO)). JLS is a Chan-Zuckerberg Biohub SF investigator. CIK and JE were funded by The Novo Nordisk Foundation, grant number NNF20CC0035580.

### Author Contributions

Conceptualization by E.B.C. and M.O.; methodology by M.O.; investigation E.B.C.; resources E.B.C., C.I.K. and J.E.; Writing –draft E.B.C.; obtaining funding, M.O., J.L.S; writing –review & editing E.B.C., M.O., J.L.S.; visualization E.B.C. and supervision M.O. and J.L.S.

### Declaration of interests

The authors declare no relevant competing interests.

Supplementary Information is available for this paper.

Correspondence and requests for materials should be addressed to Justin Sonnenburg.

Reprints and permissions information is available at www.nature.com/reprints.

## Methods

### Fermented Sample Preparation

Time course samples of kefir, curtido, kimchi, viili, and sauerkraut were prepared between 2021-2022. Milk kefir was fermented as previously described^42^. Briefly, one liter of goat milk was inoculated with 60g of kefir grains and fermented for 60 hours at room temperature. Curtido was prepared with 25% carrots, 15% onions, 4% green onions, 4% jalapenos, and 52% green cabbage by weight, all sliced with 2% salt added. After sitting for 30 minutes, curtido was jarred and fermented for 7 days before moving to the fridge for longer term storage. Both red and green sauerkraut were prepared by thinly slicing red or common (green) cabbage, adding 2% salt by weight. The initial timepoint sample was collected before letting the salted cabbage sit for 1 hour prior to jarring. For preparation of the “seeded sauerkraut,” green cabbage was prepared as described. However, just prior to jarring, 5% of cabbage by weight was removed and replaced with equal weight of a commercial common cabbage sauerkraut prepared with only cabbage and salt. Sauerkrauts were fermented for 21 days at room temperature in a jar with an airlock, prior to refrigeration. Kimchi was prepared using the standardized method obtained from the World Institute of Kimchi^53^ of 10% Korean red pepper flakes, 6% garlic, 2% sugar, 6% fish sauce, and 6% napa cabbage by weight. Napa cabbage was washed and salted with 4% salt by weight prior to rinsing and mixing with paste and jarring. Kimchi was fermented in a self-venting crock for 14 days at room temperature before being moved to 4C for long term fermentation up to 60 days. Viili was prepared by mixing 500mL of cow milk with the viili starter, and incubated at room temperature for 48 hours. All samples were collected pre-inoculation and at set time-points before being immediately frozen on dry ice and stored at −80C prior to analysis. Superior qu was prepared in Boulder, Colorado, and sampled in Liberty, Tennessee, USA, based on a recipe from Qinmin Yaoshu dating to 540 CE. For the sample preparation, equal parts of wheat berries were steamed, roasted, or ground raw and mixed with water to form a paste, formed into a stiff paste and formed into cakes, and left to dry on a mat. Every 7 days, cakes were flipped; on day 7 cakes were stacked and placed in a jar with a lid for 7 days, and finally hung until completely dry.

### Sample Collection

Fermented food samples were collected between 2021-2023, and sample source was reported (Supplementary Table 1). All samples were either collected directly into Zymo Research DNA/RNA Shield Lysis Tubes, or frozen on dry ice immediately upon collection and stored in −80C prior to DNA extraction.

### Library Preparation and Sequencing

Shotgun metagenome sequencing was performed on DNA extracted using ZymoBIOTICS DNA Miniprep kit. For samples EBC_096 through EBC_114, libraries were prepared and sequenced as previously described on a NovaSeq 6000 using S4 flow cells at Chan Zuckerberg Biohub (San Francisco, CA, USA)^36^. All other sample libraries were prepared using Roche KAPA HyperPlus and 96-UDI plates following manufacturer’s instructions. Pair-end sequencing (2×150bp) was performed using Illumina shotgun sequencing at the UC San Diego IGM Genomics Center utilizing an Illumina NovaSeq 6000 that was purchased with funding from a National Institutes of Health SIG grant (#S10 OD026929), In total, 819.1 Gbp of raw sequencing was generated.

### Metagenome quality control and assembly

Raw sequencing was demultiplexed and processed using BBtools (BBMap - Bushnell B. - sourceforge.net/projects/bbmap/), including marking of exact duplicate reads (clumpify), trimming of adapters and low-quality bases (bbduk; trimq = 16, minlen=55), duplicate reads were removed, and finally mapped against the human genome (hg19) to remove any human reads while maintaining broad eukaryote regions. FastQC (http://www.bioinformatics.bbsrc.ac.uk/projects/fastqc) was used for read quality. Metagenome assembly was performed using MegaHIT v1.2.9^54^ and binned using MetaBAT2 v2.15^55^. In total, 1186 novel bins assembled from the 90 samples (Fig. 1c) were included in the input dataset beta, described below, generating 7.21 Gbp of read pairs.

### Prokaryote database development

Eukaryotic assemblies were eliminated as a first step of developing the database consisting of bacterial and archaeal genomes. EukRep-0.6.7^56^ was used across all assemblies to identify potential eukaryotes. Genomes with >50% eukaryote assignment and a total eukaryotic genome length of >6Mbp were assigned as “eukaryotic”. dRep v3.4.5^57^ was employed on all assemblies that did not meet the criteria, assumed prokaryotic to determine species-level groups. Species groupings were based on ≥ 95% average nucleotide identity (ANI). A quality threshold was set at >50% completeness and <10% contamination for the identification of “medium” quality genomes^58^. dRep clustering identified 586 unique representative genomes. All genomes were concatenated into a.fasta file, and a scaffold was made using parse_stb.py (from dRep v3.4.5^57^) for the final MiFoDB_beta_v1_prok database. Taxonomic classification was assigned using gtdbtkv2.3.0^59^, and identification of protein-coding genes was performed using Prodigal 2.6.3^60^. GtoTree (v1.7.00)^61^ was used to make a phylogenetic tree with bacterial and archaeal gene sets, and visualized using iTol^62^ with taxonomy provided by gtdbtk, using all default settings.

### Eukaryote database development

Assemblies that met criteria for “eukaryote” assignment (>50% eukaryote assignment and >6Mbp genome length) using EukRep-0.6.7^56^ were then scored for completeness and contamination using eukCC ver_1.1^63^. Representative genomes were identified using dRep v3.4.5, with a completeness threshold >50% (-comp 50). As the contamination score of a number of NCBI reference eukaryotic genomes was >10% (potentially due to polyploidy), contamination weight was set to 0 (--contamination_weight 0). dRep clustering identified 87 unique representative genomes. Protein alignment using DIAMOND^64^ was performed against eukCC results and compared against UniRef100^65^ with a max e-value of 0.0001. Taxonomic identification of genomes without assigned taxonomy from ODFM or NCBI was performed using tRep (https://github.com/MrOlm/tRep/tree/master/bin). tRep results annotated five assembled genomes as likely substrate genomes and unrelated to microbial fermentation (Supplementary Table 4). These genomes were filtered out, and the final 82 genomes were used in the development of the final MiFoDB_beta_v1_euk database scaffold and fasta file.

### Substrate database development

For MiFoDB_sub, reference genomes from common fermented food substrates, including *Brassica oleracea* var. *oleracea* (wild cabbage plants), *Bos taurus* (cow), *Capra hircus* (goat), *Vitis vinifera* (wine grapes), *Glycine max* (soybean), *Oryza sativa* (rice), and *Triticum aestivum* (common wheat) were downloaded directly from NCBI. Genomes were concatenated into one .fasta file and scaffold file (parse_stb.py) to make MiFoDB_beta_v1_sub.

A concatenated fasta and scaffold-to-bin file was generated for all species-representative prokaryote genomes, eukaryotic microbe genomes, and substrate genomes. Files are available on Zenodo (https://zenodo.org/records/10870254).

### *Metagenome mapping and* diversity characterization

Reads from all trimmed fastq metagenomes were profiled using inStrain profile (inStrain v1.8.0^25^). Relative abundance was calculated based on # reads mapping to a genome/ total # reads in total metagenome. Shannon diversity was calculated based on relative abundance. For detailed information on profiling using MiFoDB visit: https://mifodb.readthedocs.io/

### Characterization of unmapped reads

After installing sylph 0.5.1^37^, reads from all trimmed fastq metagenomes were profiled against the GTDB-R214 based c200 database, allowing for higher sensitivity for low-abundance genomes. Resulting .tsv output was used for downstream analysis.

### Strain sharing analysis

Strain tracking was performed using inStrain compare (inStrain v1.8.0^25^). Output included a distance matrix for each species with popANI values used to cluster individual strains using “average” hierarchical clustering with a threshold of 99.999% popANI. Strains shared between sample pairs were calculated based on this ANI.

### Untargeted kefir metabolomics

LC-MS was performed as previously described^26^. Briefly, milk kefir was diluted at 1:10 in HPLC grade water and spun at 300xg for 5min. The supernatant was used for downstream processing, as previously described (Han and Van Treuren, 2021). For metabolite extraction, 100% LC-MS grade methanol containing internal standard was added (1:4 v/v). Next, samples were incubated for 5 minutes at room temperature followed by centrifugation at 5000g for 10 minutes to precipitate metabolites. Samples were transferred and dried under air to evaporate the solvent, then reconstituted in a reconstitution buffer (50:50 methanol:water v/v) with internal standards. Samples were then analyzed on the LC-MS instrument Agilent Q-TOF 6545 using reverse phase c18 positive mode, reverse phase c18 negative mode, and HILIC negative mode. Compound annotation was performed using MSDIAL (v 3.82) and an authentic standard reference library. Area under the curve for each metabolite was normalized using the sum of the internal standards in each sample. For annotated metabolites identified across multiple modalities, only the normalized peak area of the mode with the highest signal to noise ratio was used in downstream analysis.

### Metabolite-strain correlation

Metabolite normalized peak area was correlated to metagenome strain abundance (# reads mapping to a genome/ total # reads in total metagenome) across all time points using the stats (version 3.6.2) package in R. Only strains identified across >50% of time points were included in the correlation analysis.

### Functional analysis

All prokaryotic genes (n=1,542,157 detected genes) previously annotated with Prodigal 2.6.3 (Hyatt et al. 2010) were profiled against 3 databases: Pfam (protein families), CARD (antibiotic resistance genes), and the CAZy database (carbohydrate-active enzymes (CAZymes)). Pfam annotations were performed using HMMer with the commands “hmmsearch --cut_ga -- domtblout --acc Pfam-A.hmm” and “cath-resolve-hits.ubuntu-20.04” against the Pfam v34.0 database. CARD annotations were performed using the command “diamond blastp -f 6 -e 0.0001 -k 1” against the v3.2.5 database. CAZy annotations were performed using the command “hmmscan --domtblout Delta.faa_vs_dbCAN_v11.dm dbCAN-HMMdb-V11.txt GenomeSet_delta.genes.faa; sh /hmmscan-parser.sh Delta.faa_vs_dbCAN_v11.dm > Delta.faa_vs_dbCAN_v11.dm.ps; cat Delta.faa_vs_dbCAN_v11.dm.ps | awk ‘$5<1e- 15&&$10>0.35’ > Delta.faa_vs_dbCAN_v11.dm.ps.stringent” with the dbCAN v11 database. CAZyme substrate annotations were based on previously-used definitions. Results were parsed using inStrain parse_annotations (inStrain v1.8.0^25^). More detailed instructions on gene annotation and parsing can be found here: https://instrain.readthedocs.io/en/latest/user_manual.html#gene-annotation.

## Data Availability

All databases are available at: https://zenodo.org/records/10870254 The raw sequencing data for this study are in preparation under NCBI Sequencing Read Archive (SRA) SubmissionID SUB14310273 and BioProjectID PRJNA1092721

## Code Availability

https://github.com/elisacaffrey/MiFoDB

