## Supplementary Note 1 for "MiFoDB, a workflow for microbial food metagenomic characterization, enables high-resolution analysis of fermented food microbial dynamics"

*A. oryzae* is a Generally Regarded As Safe (GRAS) filamentous fungus used in production of rice and soy based ferments like miso, soy sauce, and amazake. Closely related *A. flavus* and *A. parasiticus* are pathogenic, known to infect seed crops and produce aflatoxin, a carcinogenic secondary metabolite[1]. A similar parallel might be drawn to *E. coli* and *Shigella*: while genetic relatedness would classify *E. coli* and *Shigella* as a single species, due to clinical significance the two microbes retain distinguished classifications, and research has focused on identifying accurate and reliable markers to distinguish the two[2]. While no *A. oryzae* strain has been reported to produce aflatoxin on rice and soy substrates, it retains an aflatoxin biosynthesis gene cluster[3].

We expanded our search for identifying markers of *A. oryzae* and *A. flavus* by including samples of various miso ferments made with rice koji inoculated with commercial *A. oryzae* as controls. Samples were mapped to *A. flavus* and *A. oryzae* RefSeq genomes, with all multi-mapped reads removed. Scaffolds in all samples mapped to *A. flavus*, including samples made with a commercial rice koji starter. As it is highly unlikely that *A. flavus* is present in samples made with koji from commercial *A. oryzae*, identification of *A. flavus* indicates a high false positive rate (Supplementary Fig. 6). In keeping with the goal of this database to provide information about microbes involved in food fermentation, MiFoDB_euk does not include *A. flavus*. In order to confirm that our samples do not contain *A. flavus,* genomes from a number of Aspergillus species including 7 different *A. flavus* references and 4 *A. oryzae* genomes were clustered with bins of interest using Mash[4]. Resulting dendrogram shows formation of two sub-clades, one containing *A. oryzae* reference genomes and fermented food sample bins (average ANI to sample bins = 0.99559931), and the other containing only *A. flavus* reference genomes (average ANI to sample bins = 0.99180935) (Supplementary Fig. 4). While the distinct clustering of the two species genomes (reference genomes and MAGS) offers support for the lack of *A. flavus* in the novel superior qu sample, it remains important to use caution when classifying *A. oryzae* and *A. flavus*, particularly in novel fermented food samples. Out of abundance of caution, coupling of MiFoDB-profiled metagenomics with HPLC methods to detect aflatoxin presence would allow for the highest degree of certainty when it comes to characterizing novel *Aspergillus*-based fermented foods.
