## Supplementary figures and images for "MiFoDB, a workflow for microbial food metagenomic characterization, enables high-resolution analysis of fermented food microbial dynamics"

### High resolution figures

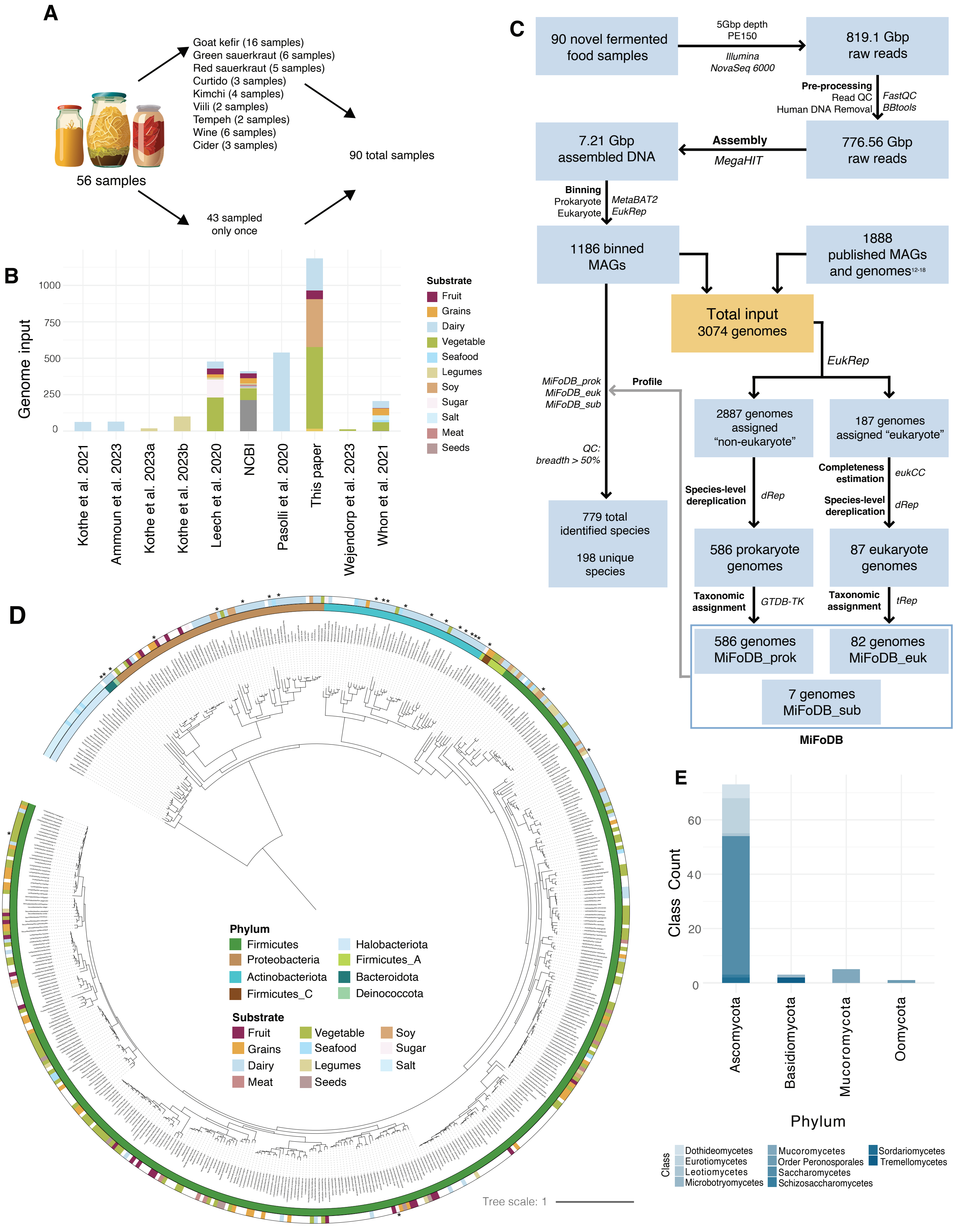

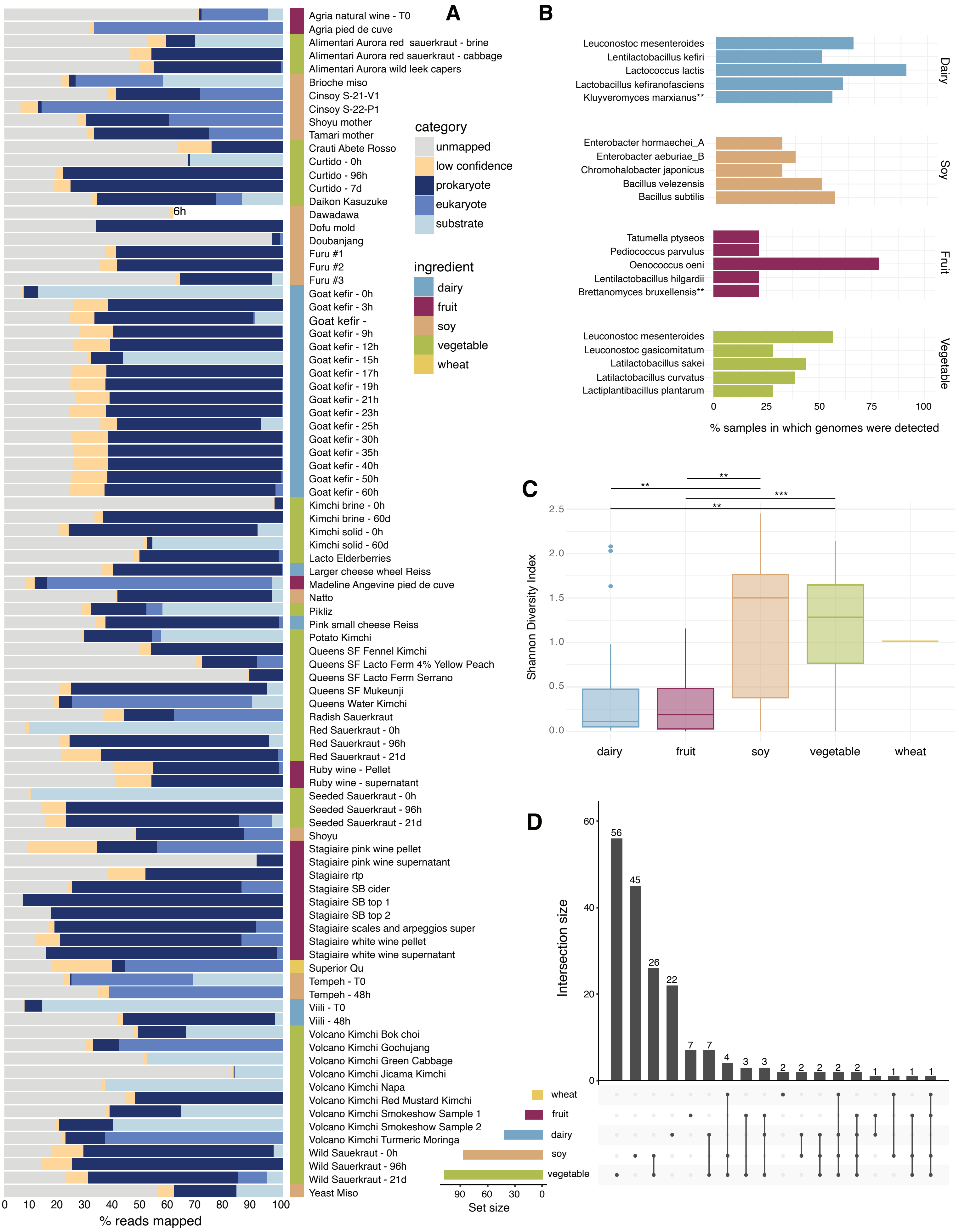

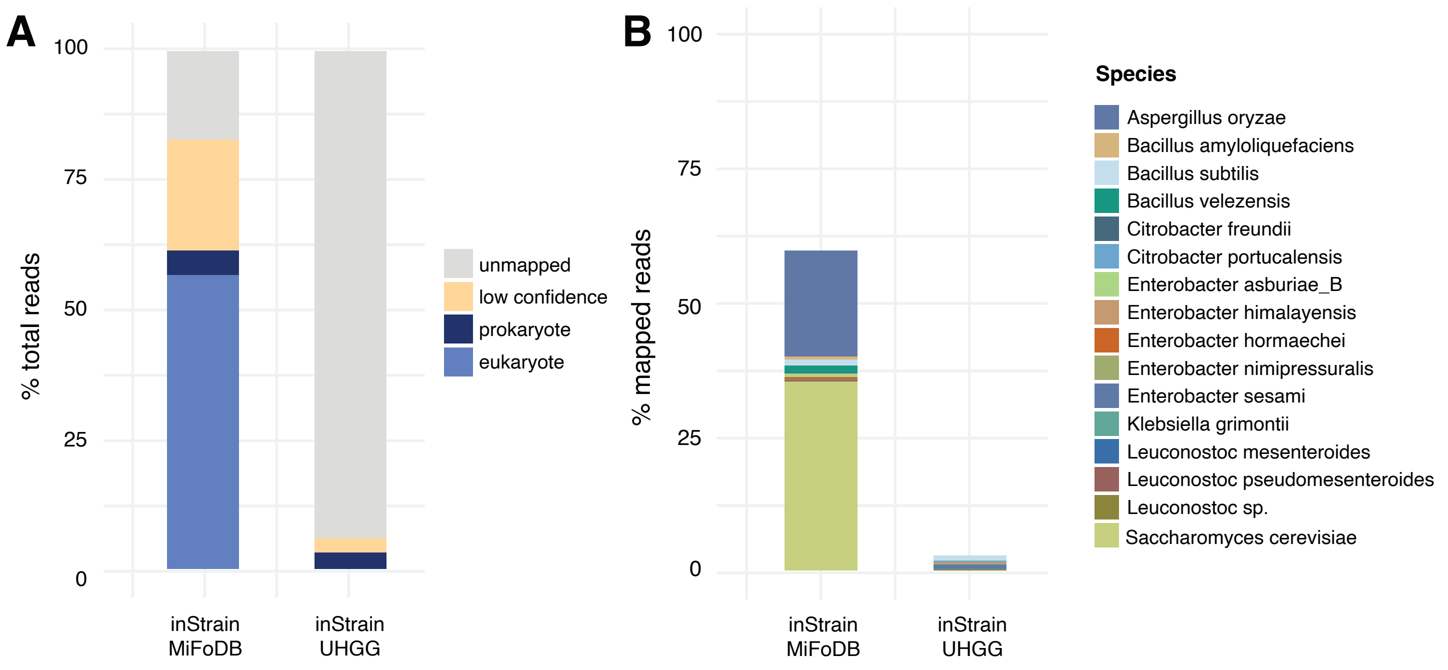

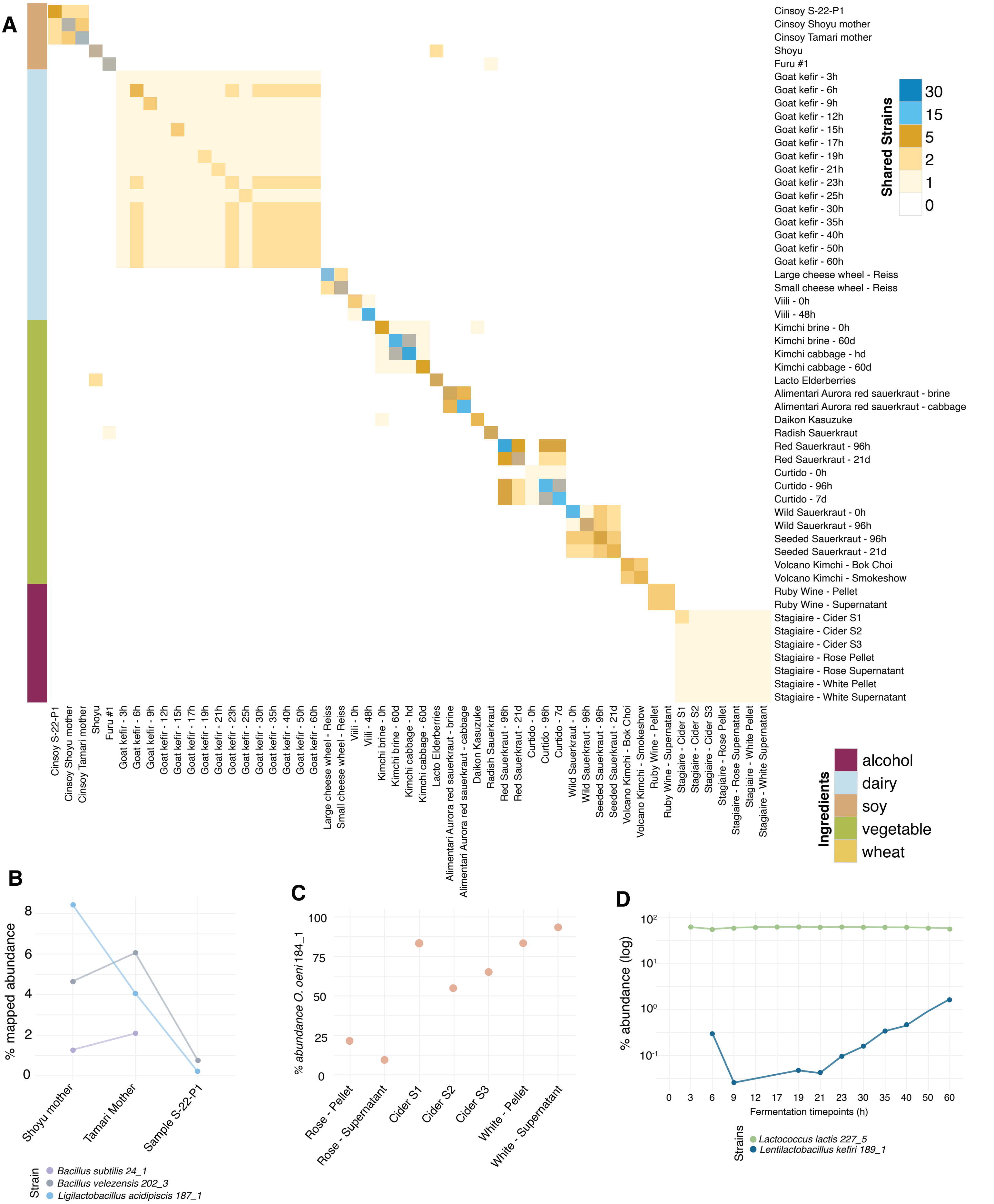

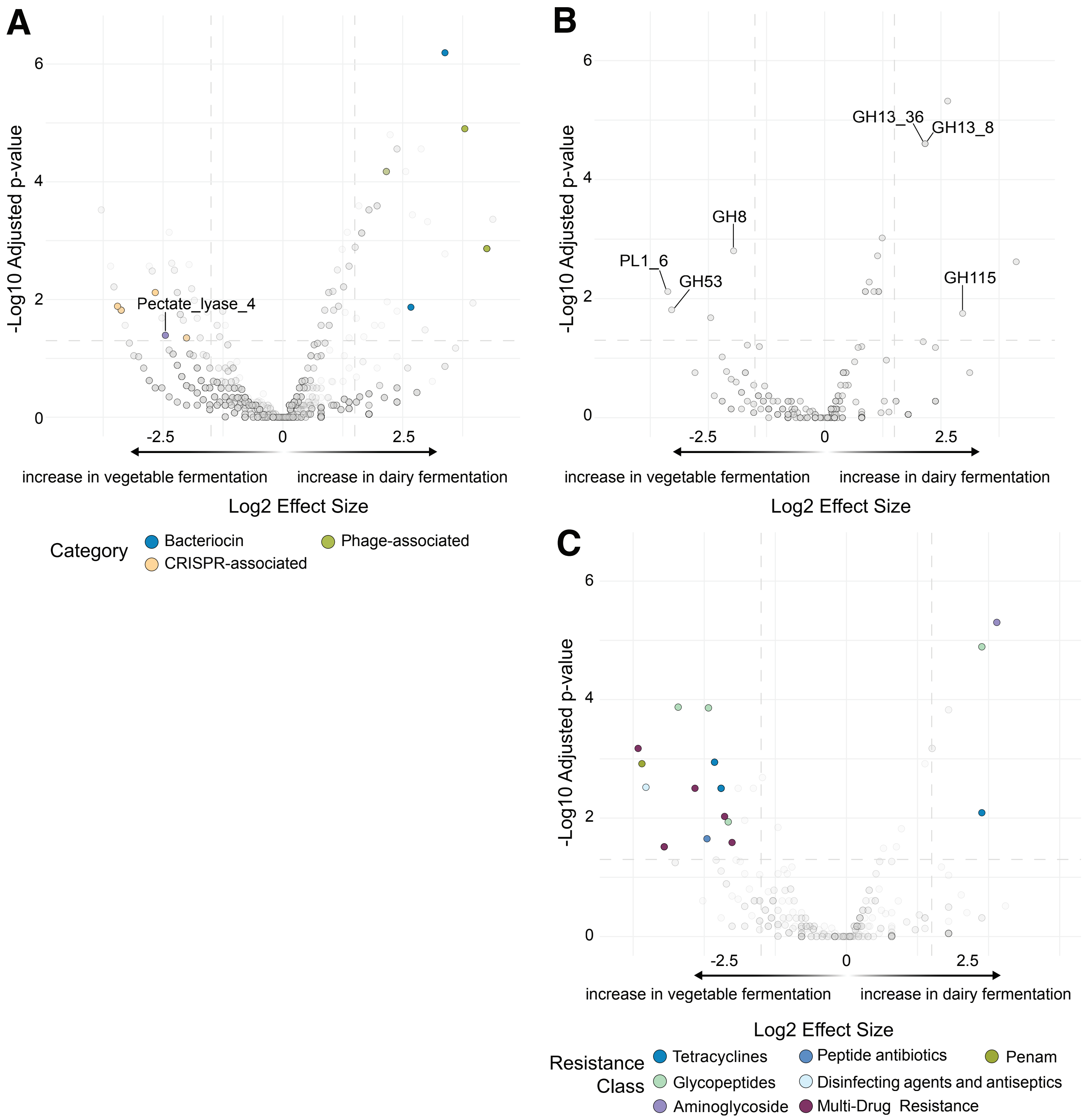
